# Early antifungal resistance prediction based on MALDI-TOF mass spectrometry and machine learning

**DOI:** 10.1101/2025.07.29.667360

**Authors:** Diane Duroux, Yiqi Yang, Janko Sattler, Michael Krauthammer, Adrian Egli

## Abstract

Antimicrobial resistance (AMR) is a significant global health threat. Recent studies have shown that combining MALDI-TOF mass spectrometry with machine learning (ML) algorithms can accelerate AMR determination. However, these efforts have predominantly focused on bacterial pathogens. Given the significant morbidity, mortality and healthcare costs associated with fungal infections and their evolving antifungal resistance to treatment, there is a need for precision medicine approaches that enable early and accurate detection of antifungal resistance.

We developed a machine-learning pipeline that integrates MALDI-TOF mass spectrometry data and drug features to predict antifungal resistance and identify spectral biomarkers for antifungal resistance. By leveraging the DRIAMS dataset, we included 658 pathogen spectra linked to 3,046 phenotypic antifungal resistance results. This dataset covered three drug classes and seven yeast species. The model was trained using categorical, phenotypic antifungal susceptibility testing results as ground truth. Our study systematically investigated the influence of various dimensionality reduction methods on MALDI-TOF mass spectra, antifungal encodings, and machine learning models using nested cross-validation to evaluate predictive performance.

We identified that applying principal component analysis to MALDI-TOF mass spectra for dimensionality reduction, and training a multi-layer perceptron yielded the highest and most stable performance for the prediction of antifungal resistance. Our method achieved an AUPRC of 0.77 across the 10 highest-performing species-drug pairs and corresponding Matthew’s correlation coefficient of 0.69. The model demonstrated the best performance for the species-drug combinations of *Candida albicans* with micafungin, *Candida parapsilosis* with fluconazole, and *Saccharomyces cerevisiae* with itraconazole and fluconazole. By comparing established species-based guidelines, susceptibility test results, and machine learning predictions, we estimated that integrating our algorithm into antifungal selection could help avoid prescriptions to likely resistant pathogens in approximately 3 out of 10 patients for whom standard guidelines recommend such treatments.

## 1 Introduction

Fungal infections are an increasingly significant global health threat, with an estimated 1.5 million deaths annually attributed to invasive fungal diseases [1]. These infections are particularly alarming for clinicians due to their high mortality rate, which often exceeds 50% [2]. Invasive fungal infections are becoming increasingly harder to treat due to the rising prevalence of antifungal resistance [3]. Unlike bacterial infections, for which a larger choice of antimicrobial treatments exists, options for combating fungal infections are far more limited. This limitation makes accurate prediction and management of antifungal resistance clinically important. As resistance continues to spread, the consequences are severe, resulting in longer hospitalizations, increased healthcare costs, and higher mortality rates [4, 5]. Given the escalating burden of fungal infections, there is an urgent need for more rapid and precise diagnostic methods to inform and optimize treatment strategies.

The determination of appropriate antifungal treatment has traditionally relied on identifying fungal pathogens using Matrix-assisted laser desorption/ionization time-of-flight mass spectrometry (MALDI-TOF MS), followed by subculture of single colonies and determination of antifungal susceptibility testing (AFST). This conventional procedure can take up to 72 hours to yield results, a delay that can be detrimental, especially for critically ill patients who require immediate and effective treatment [6].

Recent advancements have expanded the capabilities of MALDI-TOF MS, demonstrating its potential not only to accurately identify bacterial and fungal species but also to provide early insights into antimicrobial resistance [7, 8, 9, 10, 11, 12, 13]. This dual functionality could significantly improve patient outcomes by enabling more timely and targeted therapeutic interventions [14, 15, 16]. The advantages of MALDI-TOF MS in clinical settings include rapid analysis times, cost-effectiveness, and minimal sample preparation requirements, making it well-suited for routine use [17]. Moreover, the integration of MALDI-TOF MS with advanced data analysis techniques, such as machine learning, has further enhanced its potential [11, 12, 13, 18]. These innovations allow for the development of predictive models that can anticipate resistance patterns based on MALDI-TOF MS, potentially increasing the speed and precision of antifungal treatment decisions. Despite the potential of MALDI-TOF MS in antimicrobial resistance prediction, significant challenges remain. While this technology has been extensively explored for bacterial infections to predict antimicrobial resistance [19, 20, 21, 22], its application to antifungal resistance remains underdeveloped. The lower prevalence of fungal infections has led to smaller datasets [18, 23], limiting the effectiveness of machine learning models that rely on large, diverse data. Additionally, most existing MALDI-TOF MS-based antifungal susceptibility approaches require incubating fungal strains with antifungal drugs for hours to detect proteomic changes [8, 24]. While informative, these methods demand extended experimental setups and are time consuming due to the incubation with the drug, reducing their clinical practicality in urgent settings. In contrast, our approach aims to leverage MALDI-TOF spectral profiles already routinely acquired in diagnostic laboratories.

Furthermore, recent research has demonstrated that antimicrobial resistance prediction using MALDI-TOF MS can significantly benefit from large-scale deep learning models, which thrive on vast amounts of data [12, 13]. However, whether these findings extend to antifungal resistance prediction remains to be explored. The advance in drug molecular representation [25, 26], i.e. encoding molecules in machine-readable formats, could also enhance performance by leveraging more efficient representations. This underscores the importance of exploring state-of-the-art machine learning approaches to push the boundaries of antifungi resistance prediction and improve the accuracy and effectiveness of these models.

The goal of this project is to address these gaps by developing a machine learning pipeline that integrates mass spectral data with drug chemical features to predict antifungal resistance and identify the peaks contributing to the prediction. Our model will produce a binary classification indicating whether a pathogen is resistant or susceptible to a specific antifungal drug (intermediate category was not observed in the dataset). To evaluate the performance of our model, we will compare its predictions against phenotypic antifungal susceptibility testing from ISO accredited diagnostic microbiology laboratories. Additionally, to assess its potential impact on treatment selection, we will compare its predictions with species-based hospital guidelines from the University Hospital Zurich. By integrating mass spectrometry data and drug molecular structure information with advanced machine learning, this project aims to assess the diagnostic performance of MALDI-TOF MS with ML against phenotypic methods for rapid AFST.

## 2 Materials and methods

The proposed method should act as a resistance warning system, designed to alert clinicians when a pathogen is likely to be resistant to the first-line antifungal treatment, which is typically chosen based on species identification and standard hospital guidelines. When the model identifies a high probability of resistance from MALDI-TOF MS data, it will prompt clinicians to consider alternative treatment strategies, such as switching to the second-line antifungal recommended in institutional protocols. Hence, in the absence of predicted resistance, the system will remain silent, allowing clinicians to continue with standard practice unimpeded. Only when resistance is strongly suspected does the model intervene, acting as an early, evidence-based signal to re-evaluate treatment.

### 2.1 Data and pre-processing

The output to predict is a binary label for each pathogen–antifungal combination, indicating whether the pathogen is resistant or susceptible to the antifungal. The associated input for each combination consists of a MALDI-TOF mass spectrum of the pathogen and the molecular representation of the corresponding antifungal. We used the DRIAMS dataset [27] (DOI 10.5061/dryad.bzkh1899q), containing the MALDI-TOF mass spectra of pathogens and associated antimicrobial susceptibility phenotypes.

MALDI-TOF mass spectra were acquired at four microbiological laboratories in Switzerland that provide routine diagnostic services for hospitals and private practices from 2015 to 2018. Only DRIAMS-A (University Hospital Basel) and DRIAMS-C (Canton Hospital Aarau) contained fungi data. The species of each mass spectrum was identified using the Microflex Biotyper Database included in the flexControl Software (Bruker Daltonics flexControl v3.4).

DRIAMS-C was excluded from the analyses as the pathogens were only associated to susceptible profiles, for any drug. Species-drug combinations with intrinsic resistance, as indicated by EUCAST guidelines, were excluded from the dataset. This only concerned *Candida krusei* and fluconazole (11 instances). Observations involving caspofungin in association with *Candida albicans, Candida glabrata, Candida krusei, Candida parapsilosis*, and *Candida tropicalis* were excluded, as EUCAST guidelines recommend inferring its susceptibility from anidulafungin and micafungin. Species-drug pairs only associated with one outcome (either resistant or susceptible) were removed as the model could not learn from them. Supplementary Fig. 1 presents an overview of data exclusion steps and associated number of pathogens mass spectra and phenotypic antifungal resistance outcomes. After pre-processing, the dataset contained 658 unique pathogens spectra with 3,046 antimicrobial resistance phenotypes, corresponding to 22 pathogen-drug combinations.

We used the spectral representation defined in Weis et al. [28], pre-processed using the R package MaldiQuant58 v1.19 [29]. Namely the measured intensity was transformed to stabilize the variance, and smoothed. An estimate of the baseline was removed, the intensity was calibrated, and the spectra were trimmed to values in a 2,000–20,000 m/z range. Intensity measurements were binned using a bin size of 3 m/z units. Hence, each sample was represented by a vector of 6,000 features describing the sample mass-to-charge ratio and associated intensity.

### 2.2 Pathogen mass spectra dimensionality reduction

Because the number of observations is smaller than the number of variables, we evaluated three dimensionality reduction techniques of the mass spectra and compared them both against each other and against a baseline model with no dimensionality reduction. The first approach involved applying principal component analysis (PCA) to mass spectra features. To determine the optimal number of principal components, we tested a range of explained variance thresholds (0.50, 0.75, 0.85, 0.95, and 0.99). Each threshold represents the proportion of the total variance in the original data that we aimed to retain. For each threshold, we identified the minimum number of components required to reach the target level of explained variance and evaluated the downstream performance accordingly. This allowed us to balance dimensionality reduction with information retention in the feature space.

The second approach applied a masked autoencoder (MAE). An autoencoder is a type of artificial neural network that learns to compress data into a smaller representation and then reconstruct it as accurately as possible. It has two parts: an encoder that reduces the data into a compact form, and a decoder that tries to rebuild the original data from that compact form. Unlike traditional autoencoders that compress and reconstruct input data in full, the MAE randomly masks portions of each spectrum (i.e., specific mass spectra bins) and learns to reconstruct the missing values. This encourages the model to learn meaningful patterns and dependencies within the data, potentially improving its ability to predict antifungal resistance. By masking different parts of each spectrum during training, each sample can be augmented multiple times, increasing the model’s exposure to variation in the data. This made it a more flexible and robust choice compared to standard autoencoders. We tested several MAE configurations, including different encoding dimensions (64, 128, 512), numbers of augmented copies per sample (5, 10), ranges for the proportion of masked bins (0.2–0.5 and 0.3–0.6), and batch sizes (128, 256).

The third approach focused on interpretability and used mutual information (MI) to identify the most relevant features for predicting whether a pathogen is resistant or susceptible to a given antifungal. Instead of creating new composite variables, as in PCA or autoencoders, this method selects the mass spectral bins (e.g., top 64, 128, or 512) that have the strongest statistical relationship with the outcome. As a result, the features used by the model remain tied to the original spectra, making them easier to interpret and potentially more useful for biological insights. We compared the performance of models trained on these three transformed datasets against those trained on the original, untransformed data.

### 2.3 Antimicrobial embeddings

Some resistance mechanisms are shared across different species, which suggests that training models on data encompassing all species and drugs may enhance knowledge transfer and improve generalizability. To test this hypothesis, we compared several modeling strategies: one model trained on data from all species and drugs together, and other models trained separately by species, by drug, or by specific species-drug combinations. To build models that can learn across multiple drugs, it is not enough to only input information about the pathogen. The input also needs to contain information about the drug itself. Including the molecular information of a drug as part of the input can help the model understand relationships between different drugs. For example, if two drugs are chemically closely related, the pathogen may react to them in similar ways. It also means the model can learn more effectively from a larger and more diverse dataset.

We evaluated two different molecular representations of antifungal drugs for computational analysis. The first approach used the well-established Molecular ACCess Systems (MACCS) keys fingerprints [30], which summarizes the presence or absence of 166 specific chemical features in each molecule. The second method involved advanced molecular embeddings based on a deep learning model called MoLFormer [31]. This model was trained to learn patterns in molecular structures by analyzing large collections of chemical formulas (SMILES strings) from public databases like ZINC and PubChem. It is based on an algorithm originally developed for language processing (a transformer encoder model using rotary positional embeddings), allowing it to capture complex chemical features and relationships. By employing both traditional molecular fingerprints (166 variables) and advanced transformer-based embeddings (768 variables), our objective was to evaluate the benefits and drawbacks of using interpretable but simple drug encodings compared to advanced but less interpretable features.

This resulted in a total of eight input configurations for model training: for the two antimicrobial encodings (MACCS and MoLFormer), we evaluated four spectra representations, namely original data, PCA on MS, masked autoencoder on MS, and mutual information feature selection.

## 2.4 Model architecture

Our research question was framed as a binary classification problem. The sample-drug pairs were categorized into resistant and susceptible classes based on EUCAST and CLSI recommendations as in Weis et al. [32].

To classify antifungal resistance, we evaluated five different machine learning algorithms for each of the eight input configurations (Fig. 1): logistic regression, support vector machines (SVM), random forests (RF), gradient boosting classifier (GBC), and multilayer perceptron (MLP). For each method, we conducted a hyperparameter grid search, systematically testing all possible combinations within a predefined set of parameter values, to identify the combination that resulted in the best classification performance. More details on the parameters can be found in the Supplementary Section 1. This selection covers a wide spectrum of modern machine learning techniques, ranging from classical statistical methods to deep learning algorithms, ensuring a strong evaluation of predictive performance across different methodologies. As a baseline, we used no dimensionality reduction for the pathogen mass spectra, represented drugs with simple MACCS fingerprints, and applied logistic regression for classification. In total, 40 machine learning configurations were evaluated, comprising 8 input configurations combined with 5 machine learning models.

**Fig. 1:**
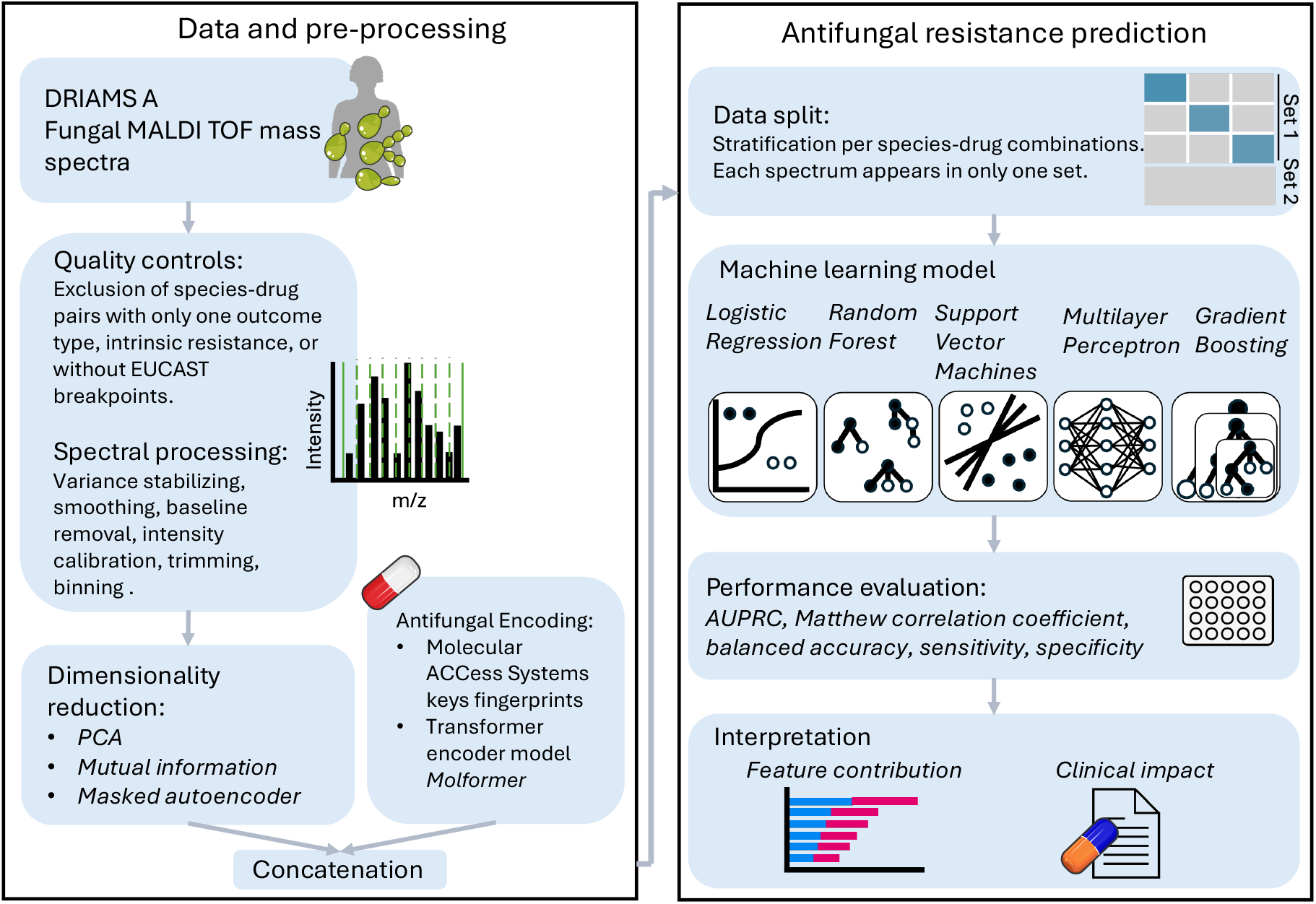
Early antifungal resistance prediction pipeline. Fungal mass spectra were selected from DRIAMS-A, a dataset derived from patient samples at University Hospital Basel. In this dataset, pathogens were cultured, their MALDI-TOF mass spectra and resistance profiles were determined, and corresponding entries were matched. Species-drug pairs associated with only one outcome (resistant or susceptible), and species with intrinsic resistance to certain antifungals were excluded. Observations involving caspofungin and *Candida spp.* were excluded too, as EUCAST guidelines recommend inferring its susceptibility from anidulafungin and micafungin. Mass spectra were pre-processed and binned following Weis et al. [32]. Three dimensionality reduction techniques for spectra and two antifungal encodings were compared. The dataset was split into Set 1 (70%) and Set 2 (30%). Nested cross-validation was performed on Set 1 to determine the optimal preprocessing methods, machine learning models, and hyperparameters. Five machine learning approaches were evaluated. After identifying the best combination, the optimal model was applied to Set 2. Model performance was assessed by comparing predictions to susceptibility test outcomes. Feature contribution analysis identified spectral peaks associated with resistance. Predictions were further compared to species-based hospital guidelines to assess the clinical impact of the tool.

### 2.5 Performance assessment

The observations were primarily divided into two sets (70% in Set 1, and 30% in Set 2). Stratification based on fungal species and drug combination was applied. We ensured that samples from the same fungal spectrum were included in either Set 1 or Set 2, but not both, to prevent information leakage and ensure unbiased performance evaluation. To identify the best configuration—i.e., the optimal combination of drug encoding, spectra representation, and machine learning model—and to perform hyperparameter tuning, we employed three-fold nested cross-validation within Set 1, with the same stratification criteria described above. Final performance are reported on Set 2. Hence, Sections 3.1 and 3.2 are based on Set 1, while the final species-drug performance results (Section 3.3, 3.4, and 3.5) are reported on a separate, held-out dataset (Set 2) that was not used during model selection.

To address the bias inherent in imbalanced datasets, we reported the Area under the precision-recall curve (AUPRC), the Matthews correlation coefficient (MCC), and the balanced accuracy. We also reported the Area Under the Receiver Operating Characteristic curve (AUROC), sensitivity, specificity, accuracy, very major error, major error, number of true positives, true negatives, false positives and false negatives. The hyperparameter search was optimized for MCC. The MCC provides a balanced measure of binary prediction performance, even in the presence of class imbalance. It ranges from -1 (total disagreement) to 1 (perfect prediction). A crucial aspect of our evaluation is the granularity of performance assessment. We focused on the species-drug combination level, as aggregated metrics can mask poor performance in specific pairs, and particularly given the over-representation of Candida albicans, which can skew overall results. Thus, we computed and reported metrics at the species-drug pair level. To identify the optimal model configuration (dimensionality reduction, drug embedding, and ML model), we averaged these pair-level metrics, aiming to select a single best configuration. Additionally, recognizing the value of strong performance in specific pairs over uniform moderate accuracy, we also reported the average performance across the top 10 highest-performing species-drug pairs for each configuration.

### 2.6 Model properties and interpretability

The analyses described in Sections 2.3, 2.4 and 2.5 allowed us to identify the best approach for predicting antifungal resistance in terms of prediction performance. Subsequently, we leveraged this optimal method to study additional capabilities of the model, which can be divided in two categories: interpretability and clinical relevance.

In terms of interpretability, our goal was to determine whether species-specific peaks drive antifungal resistance. We studied this questions through feature importance analyzes. Specifically, we identified the 500 spectral features that contributed the most to PCA and trained an MLP model on Set 1 using these features. To quantify the contributions of spectral variables to antifungal resistance, we computed SHAP values [33]. For each species-drug combination, the overall importance of a feature was determined by averaging the absolute SHAP values across correctly classified samples in Set 2. From this, we selected the top 5 features. Because we created bins of size 3 for analysis, all detected m/z should be considered as ±1. For example, if feature 3330 is reported, the corresponding m/z ratio could be 3329, 3330, or 3331.

To evaluate the clinical relevance of our model, we assessed both the risks of incorrect antifungal resistance predictions and its potential benefits over species-based guidelines for antifungal selection. Specifically, we examined whether a patient would still receive an antifungal associated with susceptibility (based on susceptibility tests) if the model incorrectly predicts resistance and how often integrating the model into treatment decisions could prevent antifungal prescriptions associated with resistance compared to species-based guidelines. To answer these questions, we compared the model’s predictions with species-based guidelines from the University Hospital Zurich (2024), which are available upon request (https://www.usz.ch/fachbereich/infektiologie/ueber-uns/bestellen-sie-unsere-antibiotikarichtlinien/).

## 3 Results

### 3.1 Combination of dimensionality reduction and neural networks give rise to the highest antifungal prediction performance

The dataset comprises fungi from seven species, including various *Candida* species and *Saccharomyces cerevisiae* (Table 1). It includes critical fungal species identified by the WHO, such as *Candida albicans* and *Candida glabrata* [34]. The dataset spans seven drugs, including representatives from the azole class and echinocandins, resulting in a total of 22 species-drug combinations.

**Table 1.** Summary of datasets. Set 1 contains 70% of the total observations, and Set 2 contains 30%.

|  | Number of<br>antifungal resistance<br>phenotypes | Number of<br>species-drug<br>combinations | Number of<br>unique pathogen<br>spectra | Number of<br>unique species | Number of<br>unique drugs | Proportion of<br>resistant<br>observations |
| --- | --- | --- | --- | --- | --- | --- |
| **Set 1** | 2123 | 22 | 460 | 7 | 7 | 0.18 |
| **Set 2** | 923 | 22 | 198 | 7 | 7 | 0.16 |

Set 1 consists of 460 pathogen spectra, while the Set 2 includes 198 spectra. Each observation in our dataset represents a pathogen-drug pair, as multiple drugs are tested on individual pathogen isolate. Consequently, Set 1 contains 2,123 antifungal resistance phenotypes, and Set 2 includes 923 phenotypes. Across the entire dataset (Set 1 and 2 combined), the number of samples per species varies widely (Fig. 2 (a)), ranging from 11 for *Candida krusei* to 300 for *Candida albicans*. Similarly, the number of observations varies by drug, with 334 samples tested on posaconazole and 518 on anidulafungin (Fig. 2 (b)). The majority of samples were collected in 2016 (212 samples), 2017 (268 samples), and 2018 (157 samples).

**Fig. 2:**
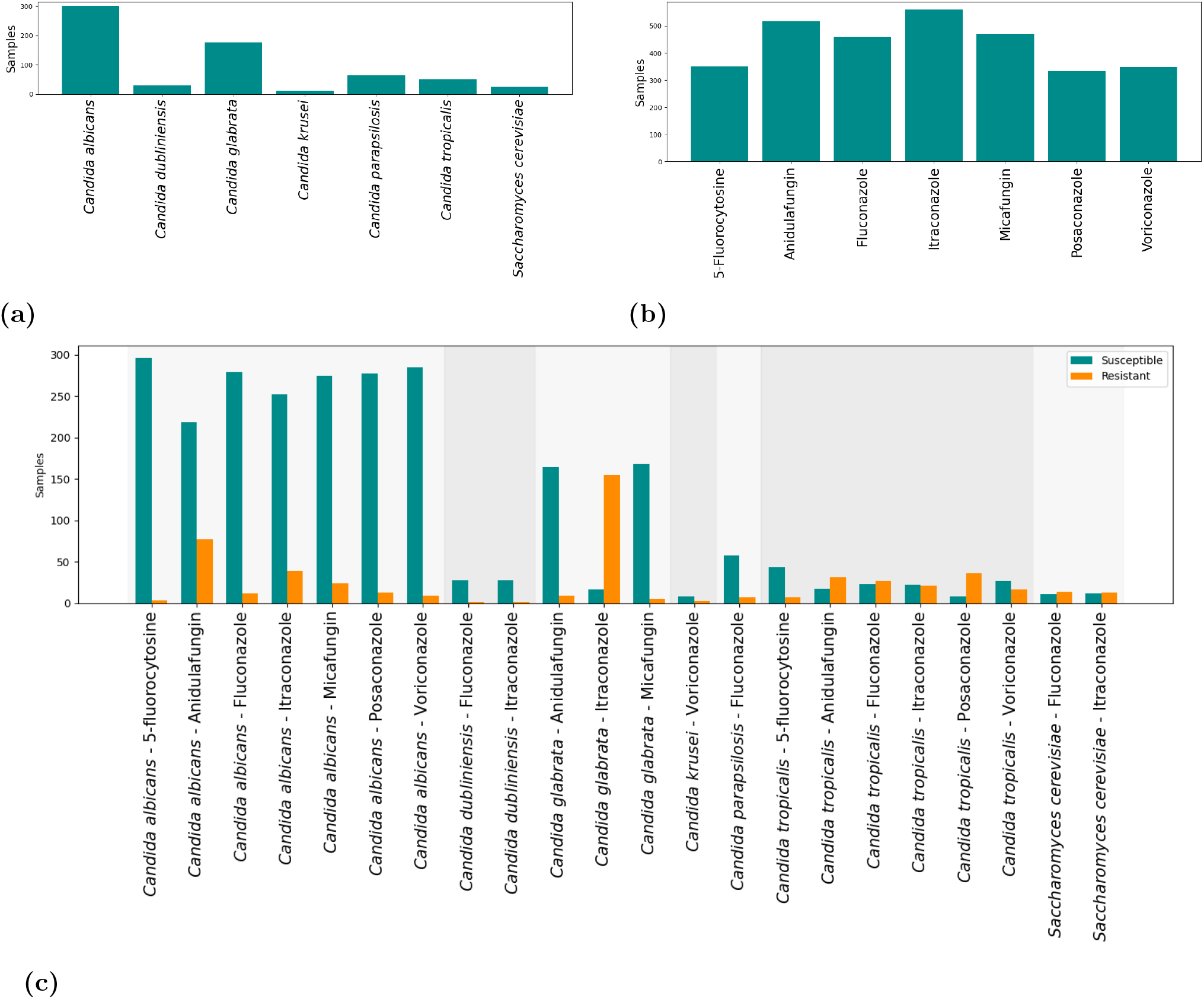
Data distribution of the fungi dataset based on (A) species, (B) drugs, and (C) species-drug pairs with response 0 for susceptibility and 1 for resistance.

Fig. 2 (c) highlights the dataset’s class imbalance, with some species-drug pairs showing significant disparity. For instance, the *Candida albicans*-5-fluorocytosine pair exhibits a stark imbalance, with 296 susceptible versus only 4 resistant outcomes. In contrast, some pairs are nearly balanced, such as *Candida tropicalis*-itraconazole, with 22 susceptible and 21 resistant outcomes. Most species-drug combinations show a higher number of susceptible cases compared to resistant ones, although the opposite is observed in specific cases, such as *Candida glabrata*-itraconazole and *Candida tropicalis*-posaconazole. This inherent imbalance necessitated the use of evaluation metrics like balanced accuracy, AUPRC, and MCC to ensure a robust assessment of model performance.

Our study systematically investigated the influence of various dimensionality reduction methods on MALDI-TOF MS, antifungal encodings, and machine learning models, on the predictive performance (Fig. 3). We found that Multilayer Perceptron (MLP) neural networks and Support Vector Machines (SVM) outperformed other models across MCC (Fig. 3), AUPRC, and balanced accuracy metrics (supplementary Fig. 2 and 3). Among the top 10 machine learning configurations out of the 40 evaluated, the MLP model appeared six times, while the SVM appeared four times. A similar trend, with MLP and SVM demonstrating superior performance, was observed when considering all species-drug pairs for MCC computation (supplementary Fig. 4).

**Fig. 3:**
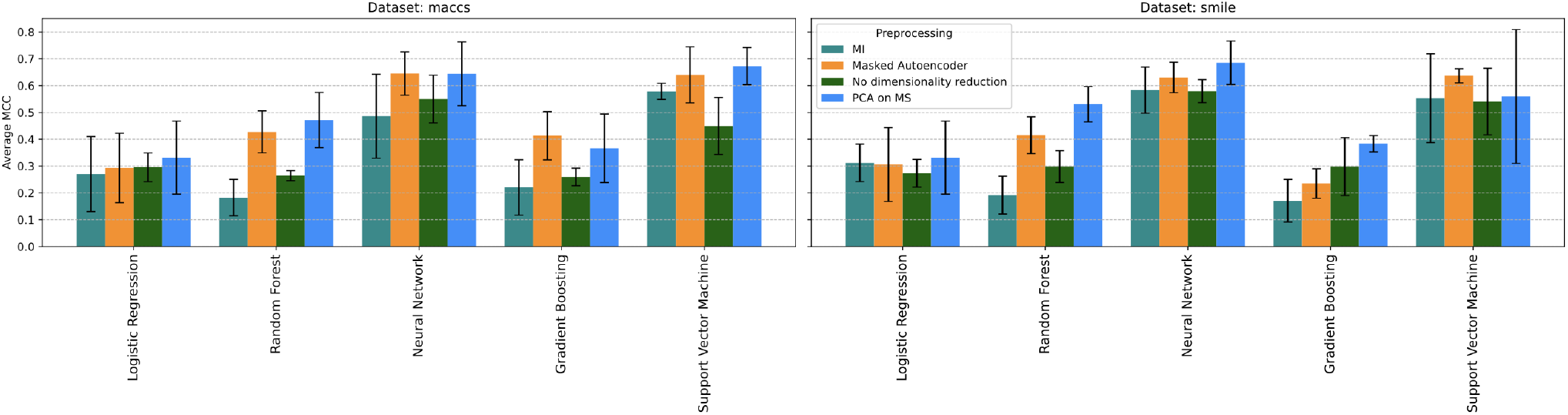
Performance evaluation (MCC) across pre-processing techniques and models for the top 10 highestperforming species-drug pairs. Error bars show the standard deviation computed via the 3-fold cross validation.

Regarding dimensionality reduction of mass spectra, PCA and MAE demonstrated superior performance compared to mutual information and the absence of dimensionality reduction. Among the top 10 configurations out of the 40 evaluated, the MAE appeared four times, the PCA three times, mutual information two times and no preprocessing one time. Across all configurations, the mean MCC values for each dimensionality reduction method of the mass spectra were as follows: mutual information (0.35), no preprocessing (0.38), MAE (0.46) and PCA on MS (0.5).

For antifungal encoding, no clear performance differences were observed between MACCS fingerprints and Molformer SMILES encodings. Across all configurations, the mean MCC values were 0.42 for MACCS and 0.43 for Molformer SMILES encodings.

The highest performance among all tested configurations was achieved using a combination of PCA for mass spectra, Molformer SMILES antifungal encodings, and the MLP model, yielding an AUPRC of 0.77 and an MCC of 0.69 across the top 10 highest-performing species-drug pairs. The best parameters included PCA with 99% variance retention and an MLP with two hidden layers (50 neurons each), a maximum of 1000 iterations, a tanh activation function, and a regularization alpha of 0.0001. This optimal machine learning configuration aligns with the overall performance patterns observed above. Therefore, we selected this configuration for the subsequent analyses.

### 3.2 Unified species-and-drug-agnostic modeling matches species-drug-specific performance and improves applicability

Incorporating drug fingerprints into the model enables the development of a unified prediction framework across all species and drugs. This approach contrasts with training separate models per species or per drug, which could allow for more targeted detection of species-specific resistance mechanisms but may sacrifice the benefits of knowledge sharing across combinations. To evaluate these strategies, we compared model performance across different configurations using nested crossvalidation on Set 1, applying PCA to mass spectra, Molformer SMILES encodings, and an MLP classifier. To promote stable and reliable evaluation, performance metrics are restricted to species-drug pairs with no more than 5% imbalance between classes (ratio of resistant to susceptible observations) and at least two resistant samples.

Our results showed minimal differences across model types. The unified model trained on all observations achieved a mean MCC of 0.49 (±0.02), close to models trained per species (0.46 ±0.06), per drug (0.47 ± 0.06), or per species-drug combination (0.49 ±0.03). While species-drug-specific models achieved higher performance on some clinically relevant pairs, for example, *Candida tropicalis* with anidulafungin and *Candida albicans* with fluconazole, they identified only one additional high-performing pair (MCC *≥* 0.04) compared to the unified model. Conversely, the unified model offers broader applicability by enabling predictions for rare or low-sample-size species-drug pairs.

Additionally, we assessed the impact of incorporating drug class into the model. Antifungal drugs were categorized into azoles, echinocandins, and 5-fluorocytosine, and encoded using one-hot encoding. Including drug class via one-hot encoding had no impact on performance, likely because this information is already embedded in the molecular representations.

Given the comparable overall performance and broader coverage, we selected the model trained on all observations for subsequent analyses. We refer to this configuration as the *unified* model.

### 3.3 Enhanced antifungal detection in *Saccharomyces cerevisiae, Candida albicans*, and *Candida parapsilosis*

All subsequent results are based on analyses performed using the separate Set 2 to ensure independent evaluation. To avoid skewed performance metrics, improve statistical validity and enhance generalizability, we report the unified model’s performance across the species-drug combinations where the maximum imbalance was 5% (either in resistance or susceptibility) and where at least two resistant samples were available, resulting in 12 pairs. AUPRC, MCC, balanced accuracy, number of true positives, true negatives, false positives and false negatives are reported in Table 2. Sensitivity, specificity, accuracy, very major error and major error are provided in Table 3.

**Table 2.** Performance metrics on the held-out Set 2, sorted by MCC, for the species-drug pairs with maximum imbalance 5% (either in resistance or susceptibility) and at least two resistant samples. *Imbalance* refers to the ratio of resistant to susceptible observations.

| Species | Drug | MCC | AUPRC | AUROC | Imbalance | TP | TN | FP | FN |
| --- | --- | --- | --- | --- | --- | --- | --- | --- | --- |
| *Saccharomyces cerevisiae* | Itraconazole | 1.00 | 1.00 | 1 | 0.63 | 5 | 3 | 0 | 0 |
| *Candida albicans* | Micafungin | 0.79 | 0.81 | 0.98 | 0.06 | 4 | 84 | 1 | 1 |
| *Saccharomyces cerevisiae* | Fluconazole | 0.65 | 0.73 | 0.5 | 0.75 | 6 | 1 | 1 | 0 |
| *Candida parapsilosis* | Fluconazole | 0.55 | 0.81 | 0.96 | 0.15 | 1 | 17 | 0 | 2 |
| *Candida tropicalis* | Posaconazole | 0.42 | 0.95 | 0.65 | 0.87 | 8 | 2 | 0 | 5 |
| *Candida tropicalis* | Voriconazole | 0.41 | 0.75 | 0.48 | 0.60 | 3 | 6 | 0 | 6 |
| *Candida albicans* | Itraconazole | 0.36 | 0.41 | 0.77 | 0.13 | 4 | 72 | 4 | 7 |
| *Candida albicans* | Anidulafungin | 0.32 | 0.54 | 0.70 | 0.26 | 10 | 56 | 9 | 13 |
| *Candida tropicalis* | Itraconazole | 0.26 | 0.87 | 0.59 | 0.73 | 6 | 3 | 1 | 5 |
| *Candida glabrata* | Itraconazole | 0.25 | 0.99 | 0.88 | 0.92 | 41 | 2 | 2 | 7 |
| *Candida tropicalis* | Fluconazole | 0.00 | 0.68 | 0.43 | 0.5 | 3 | 4 | 3 | 4 |
| *Candida tropicalis* | Anidulafungin | -0.58 | 0.60 | 0.37 | 0.6 | 4 | 0 | 6 | 5 |

**Table 3.** Balanced accuracy, sensitivity, specifiticy, accuracy, very major error, and major error for species-drug combinations with more than 2 resistant samples in Set 2 and maximum imbalance 5% (either in resistance or susceptibility).

| Species | Drug | Balanced acc | Sensitivity | Specificity | Accuracy | VME | ME |
| --- | --- | --- | --- | --- | --- | --- | --- |
| *Saccharomyces cerevisiae* | itraconazole | 1.00 | 1.0000 | 1.0000 | 1.0000 | 0.0000 | 0.0000 |
| *Candida albicans* | micafungin | 0.89 | 0.8000 | 0.9882 | 0.9778 | 0.2000 | 0.0118 |
| *Saccharomyces cerevisiae* | fluconazole | 0.75 | 1.0000 | 0.5000 | 0.8750 | 0.0000 | 0.5000 |
| *Candida parapsilosis* | fluconazole | 0.67 | 0.3333 | 1.0000 | 0.9000 | 0.6667 | 0.0000 |
| *Candida tropicalis* | posaconazole | 0.81 | 0.6154 | 1.0000 | 0.6667 | 0.3846 | 0.0000 |
| *Candida tropicalis* | voriconazole | 0.67 | 0.3333 | 1.0000 | 0.6000 | 0.6667 | 0.0000 |
| *Candida albicans* | itraconazole | 0.66 | 0.3636 | 0.9474 | 0.8736 | 0.6364 | 0.0526 |
| *Candida albicans* | anidulafungin | 0.65 | 0.4348 | 0.8615 | 0.7500 | 0.5652 | 0.1385 |
| *Candida tropicalis* | itraconazole | 0.65 | 0.5455 | 0.7500 | 0.6000 | 0.4545 | 0.2500 |
| *Candida glabrata* | itraconazole | 0.68 | 0.8542 | 0.5000 | 0.8269 | 0.1458 | 0.5000 |
| *Candida tropicalis* | fluconazole | 0.5 | 0.4286 | 0.5714 | 0.5000 | 0.5714 | 0.4286 |
| *Candida tropicalis* | anidulafungin | 0.22 | 0.4444 | 0.0000 | 0.2667 | 0.5556 | NaN |

The top-performing species-drug pairs in Set 2 are *Saccharomyces cerevisiae*–itraconazole, which achieved perfect predictions, *Candida albicans*–micafungin (MCC 0.79), *Saccharomyces cerevisiae*–fluconazole (MCC 0.65), and *Candida parapsilosis*–fluconazole (MCC 0.55). In terms of species, the model performed best on *Saccharomyces cerevisiae*, achieving an average MCC of 0.83 across drugs, followed by *Candida parapsilosis* (MCC 0.55) and *Candida albicans* (MCC 0.49). Among antifungals, the model performed best with micafungin (average MCC 0.79 across species), followed by itraconazole (MCC 0.38) and fluconazole (0.24). These findings highlight the model’s ability to generalize across a diverse range of species and drugs.

We observed significant performance variability across species-drug pairs and even between different drugs for the same species. For instance, while the model achieved strong results for *Candida albicans* with micafungin, it performed poorly when paired with posaconazole or voriconazole.

Note that close results are obtained with the species-drug specific models (Supplementary Table 1). For *Candida tropicalis*-anidulafungin, we obtained a negative MCC, indicating that the model performed worse than random guessing for this species-drug pair. This result may be due to the limited representation of *C. tropicalis*–anidulafungin samples in the training set, which could have prevented the model from learning meaningful resistance patterns specific to this combination. Alternatively, it may reflect intrinsic limitations in the biological signal captured by MALDI-TOF MS for detecting resistance to anidulafungin in *C. tropicalis*, suggesting that the spectral features available may not sufficiently differentiate between resistant and susceptible isolates for this particular case.

Since the model is intended as a resistance warning system, it is designed to have no impact on clinical workflow when no resistance is detected. However, when the model flags resistance, it is critical, as it may change treatment decisions. Because only resistance predictions carry the potential to influence clinical decisions, it is essential that these alerts are cautious. False positives in this context could lead to unnecessary deviation from effective therapies. We prioritize high specificity for resistance detection, ensuring that recommendations to alter treatment are reliable. Consequently, in our application, controlling major errors (false predictions of resistance) is especially important, as these are the only outcomes that could alter clinical decisions. Thus, a low major error rate is prioritized even above minimizing very major errors.

Overall, our analysis demonstrated the model’s effectiveness in predicting antifungal resistance across various *Candida* and *Saccharomyces* species-drug pairs, with particularly strong performance for *Saccharomyces cerevisiae* resistance to itraconazole and fluconazole, and *Candida albicans* resistance to micafungin.

### 3.4 Identification of spectral peaks driving antifungal resistance

The model performs best on *Candida albicans, parapsilosis, tropicalis* and *Saccharomyces cerevisiae*. Figure 4 (a) shows the difference in average mass spectra between *Candida albicans* isolates that are resistant (blue) and susceptible (orange) to micafungin. The minimum peak height value considered in this analysis is 4.3 *×* 10^*−4*^. The SHAP values, which indicate the contribution of each m/z feature to the classification model, were computed from 87 correctly classified observations. Among these, features corresponding to m/z 2076, 3474, 6909, 6921 and 6954 (*±*1), emerged as the most influential in distinguishing resistant from susceptible isolates, all exhibiting higher intensity in resistant strains. These peaks may represent potential biomarkers linked to micafungin resistance in *C. albicans*.

**Fig. 4:**
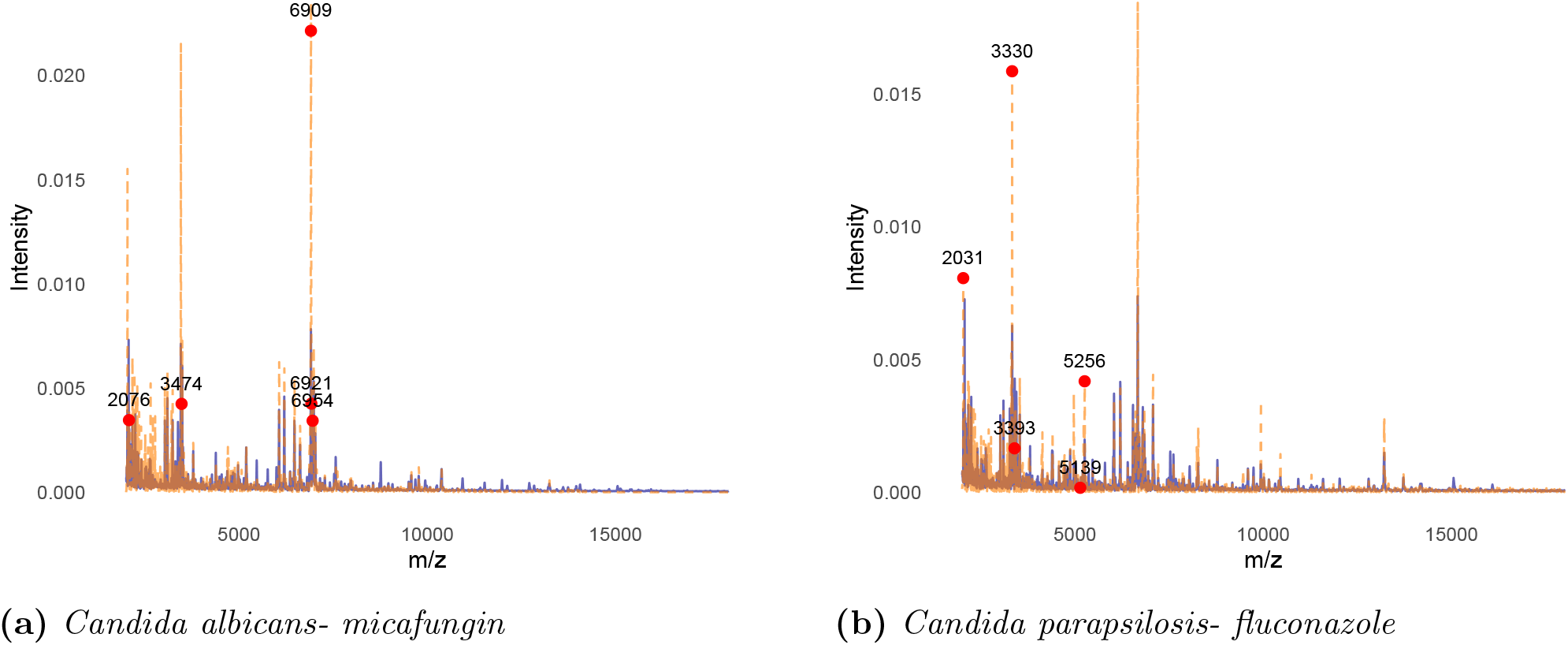
Average mass spectra for susceptible (blue) and resistant (dashed orange) pathogens. Red dots indicate the top-5 peaks that contribute most to AMR prediction (Section 2.6).

Figure 4 (b) similarly illustrates the average mass spectra differences for *Candida parapsilosis* isolates resistant (blue) and susceptible (orange) to fluconazole. The min peak height value was set to 4.5 *×* 10^*−4*^, and SHAP values were computed using 18 correctly classified samples. The five most discriminative m/z features identified in this analysis were 2031, 3330, 3393, 5139, and 5256 (*±*1). Additional SHAP-based spectral comparisons for *Saccharomyces cerevisiae* with itraconazole and fluconazole, as well as *Candida tropicalis* with posaconazole, are provided in the supplementary materials. Overall, this approach has enabled the identification of specific mass spectral biomarkers associated with antifungal resistance.

### 3.5 Estimated clinical impact: reduced selection of antifungals associated with resistance

In clinical practice, not all predictions hold the same value, as certain antifungal drugs are rarely used for particular species. To evaluate the utility of our ML approach, we determine the frequency at which a recommended antifungals would have failed according to the antifungal susceptibility tests in the DRIAMS dataset. We determine the percentage of incorrect recommendations using species-based guidelines, which define the expected activity of antifungal agents against specific fungal species. For each species, the guidelines indicate whether a given agent is considered: *generally effective and recommended for therapy, generally effective, variably effective*, associated with a *high probability of resistance*, or if *no or insufficient data* are available. We limit our analyses to the 7 drugs and 6 species present in both the DRIAMS data and for which we also have guideline recommendations (184 samples in Set 2). These include only *Candida species*. For many species, multiple antifungals are equally recommended as the top choices (i.e., *generally effective and recommended for therapy*). Since no additional patient information is available in the dataset, we hypothesize that physicians would choose randomly among them. For example, if a hospital recommends three drugs equally for a species and two of them are effective according to the susceptibility tests, we consider the hospital’s recommendation to be 66% correct for the pathogen investigated. In Set 2, using the hospital species-based recommendations, 9% of the pathogens have at least a 25% risk of being resistant to the received drug according to susceptibility tests. These cases involved *Candida albicans, Candida dubliniensis, Candida glabrata, Candida parapsilosis*, and *Candida tropicalis*. The antifungals associated with these cases are fluconazole, anidulafungin, and caspofungin. Importantly, antifungal susceptibility tests indicated that an antifungal associated with susceptibility could have been provided in 100% of these cases.

We applied the unified model (Section 3.1) to the 9% of pathogens with at least a 25% risk of resistance in Set 2. These predictions are particularly challenging as species-based guidelines do not apply, i.e., top antifungals are generally expected to be highly effective based on guidelines but fail against the specific tested pathogen according to the susceptibility tests. Within this subset, the model successfully identified resistance in 29% of cases. This indicates that among patients for whom species-based guidelines recommend antifungals to which their pathogens are likely resistant (based on susceptibility tests), the ML model could have flagged resistance earlier than conventional susceptibility testing in approximately 3 out of 10 cases (29%).

We also assessed how often the model incorrectly predicted resistance for pathogens that were actually susceptible to an antifungal and the potential impact of such errors. The model misclassified 3% of cases as resistant. However, in every instance, at least one alternative top-ranked antifungal was still predicted correctly susceptible by the ML model, ensuring that physicians could still select an antifungal likely to be effective.

## 4 Discussion

To assess the potential for early detection of antifungal resistance using MALDI-TOF MS, we explored a range of machine learning architectures and compared findings to phenotypical results. Our experiments showed that using more complex models to encode drugs molecular information did not consistently lead to better model performance. While the top-performing configuration incorporated the transformer-based Molformer-derived drug embeddings, overall results were comparable between the Molformer encodings and simpler MACCS fingerprints. Also, the more complex dimensionality reduction method for mass spectra, Masked Autoencoder (MAE), did not yield performance gains compared to simpler approaches like PCA. Several factors could explain why these more sophisticated methods failed to outperform simpler alternatives. One reason can be overfitting, as deep learning models like MAE typically require large datasets to generalize effectively. In our case, the base dataset (without dimensionality reduction) comprises 3,046 observations, with 6,000 mass spectra features and between 166 and 768 drug-related features. Additionally, in this project, we used Molformer model checkpoints pre-trained on only 10% of a larger dataset—approximately 100 million molecules, with 10% sourced from ZINC and another 10% from PubChem, originally used for training Molformer-XL. Gaining access to a Molformer model trained on the full dataset, rather than just a 10% subset, could enhance performance by providing richer and more comprehensive molecular encodings.

Most MALDI-TOF-based susceptibility testing protocols differ from our approach as they require an incubation period of at least three hours with varying antifungal concentrations. To assess the effectiveness of our direct prediction approach, we compared its performance with results reported for these incubation-based methods. Focusing on species-drug pairs in Table 2, we identified published performance metrics for three pairs. For *Candida tropicalis*-fluconazole, Saracli et al. [10] observed a very major error (VME; false susceptible/total resistant) of 0.14 (1/7) and a major error (ME; false resistant/total susceptible) of 0.43 (12/28). Our model produced a higher VME of 0.57 (4/7) and a close ME of 0.42 (3/7). For *Candida tropicalis*-posaconazole, Saracli et al. reported a VME of 0.5 (2/4) and a ME of 0.18 (6/33), while our model yielded improved VME of 0.38 (5/13) and ME of 0 (0/2). For *Candida tropicalis*-voriconazole, Saracli et al. found a VME of 0.4 (2/5) and an ME of 0.36 (10/28), whereas our model produced a higher VME of 0.67 (6/9) but a ME of 0 (0/3). Thus, our method consistently achieves a lower major error rate, demonstrating greater precision in avoiding false resistance predictions. However, it tends to underpredict resistance, resulting in higher very major errors in some cases. This balance aligns with our *warning detection* approach: if resistance goes undetected, the clinician’s workflow remains unchanged. However, when the model flags resistance, it is critical, as it may change treatment decisions. Compared to MALDI-TOF-based susceptibility testing, a key advantage of our approach is that it eliminates the need for additional laboratory experiments, reducing both processing time and costs, making it more suitable for routine clinical use.

Integrating early antifungal resistance prediction based on MALDI-TOF mass spectra into routine hospital diagnostics presents both exciting opportunities and practical challenges. The vision is to enhance existing MALDI-TOF workflows—already widely used for rapid microbial identification, by embedding machine learning models capable of simultaneously predicting resistance profiles. However, several hurdles must be addressed before clinical implementation. These include standardizing data preprocessing across different MALDI-TOF instruments, laboratory protocols, and software environments, as well as ensuring robust model validation across diverse patient populations, hospitals, geographic regions, and time periods. Seamless integration into laboratory information systems is also essential, ideally with an intuitive, clinician-friendly dashboard that fits naturally into existing diagnostic workflows. Additionally, regulatory approval, clinician trust in AI-assisted tools, and clear interpretability of model outputs will be critical for adoption.

Future evaluations will be necessary to further validate our findings and assess the real-world impact of our approach. Since our results are based on data from the University Hospital Basel, validating the model’s performance on an external dataset would strengthen confidence in its generalizability. Additionally, the spectral m/z units identified as contributors to resistance should also be experimentally validated, for instance, by integrating sequencing data to confirm their biological relevance. This is particularly important given that resistance mechanisms are often complex and context-dependent, requiring validation beyond computational inference. Incorporating genomic data in future work would allow for peak annotation and biological interpretation. Moreover, our approach uses binary resistance/susceptibility labels, whereas working with continuous outcomes, such as minimum inhibitory concentrations (MICs), could provide a more nuanced and informative representation of antifungal resistance. Finally, conducting a prospective multi-center randomized controlled trial will help validate our findings and assess whether our model would influenced treatment decisions in a clinical setting, including its potential impact on antifungal de-escalation strategies. These future steps will be essential in translating our findings into practical clinical applications.

## 5 Data and code availability

The code necessary to reproduce this article’s results and analyses is available on GitHub at https://github.com/DianeDuroux/AntifungalResistancePrediction/

## 8 Acknowledgments

We are grateful to Prof. Dr. Dr. med. Silvio Brugger for valuable advice and interesting discussions. Some images are adapted from Servier Medical Art (https://smart.servier.com/), licensed under CC BY 4.0 (https://creativecommons.org/licenses/by/4.0/).

