## Supplementary material for "Early antifungal resistance prediction based on MALDI-TOF mass spectrometry and machine learning"

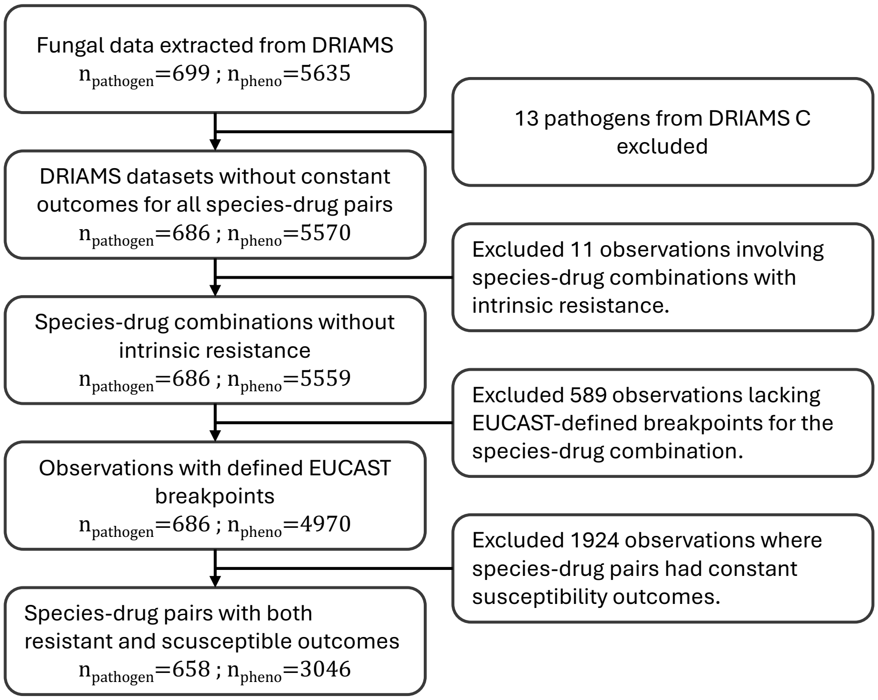


**Fig. 1:** Overview of data exclusion steps during preprocessing. $n_{pathogen}$ is the number of unique pathogen mass spectra in the dataset and $n_{pheno}$ is the number of associated phenotypic antifungal resistance results. An observation is a pathogen-drug pair.

### 1 Machine learning approaches parameters

Logistic Regression (LR) was implemented with `max_iter` set to 2000 to ensure convergence during optimization. We evaluated the model using C values of 0.1, 1, and 10, with both L1 and L2 regularization penalties.

Support Vector Machines (SVM) with a Radial Basis Function (RBF) kernel were employed, using C values of 0.1, 1, and 10 to balance error minimization and model complexity. The `probability` parameter was set to `True` to enable class probability estimation.

For Random Forest (RF), we configured the model with 50, 100, and 200 trees (`n_estimators`) and tested maximum depths of None, 10, and 20 (`max_depth`). All other parameters were kept at

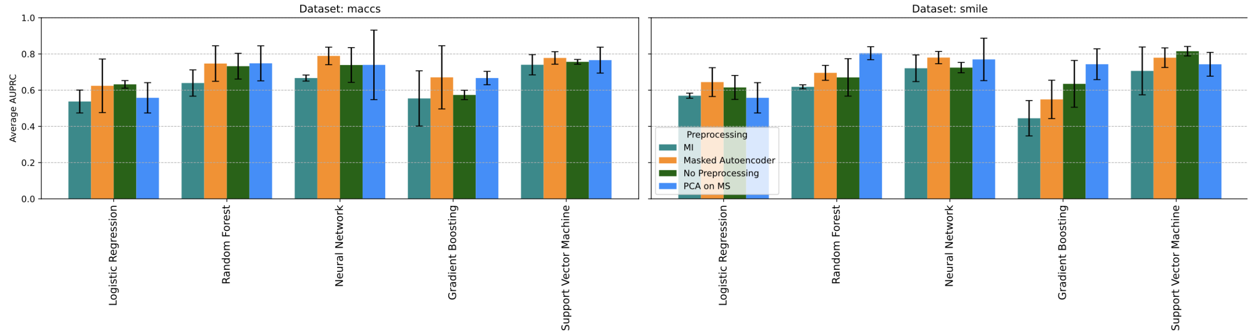


**Fig. 2:** AUPCR across pre-processing techniques and models for the top 10 highest-performing species-drug pairs. Error bars show the standard deviation computed via the 3-fold cross validation.

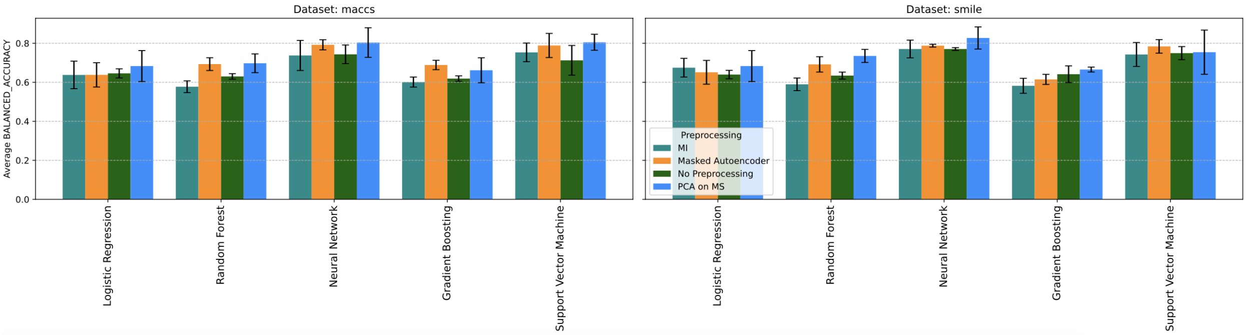


**Fig. 3:** Balanced accuracy across pre-processing techniques and models for the top 10 highest-performing species-drug pairs. Error bars show the standard deviation computed via the 3-fold cross validation.

their default values as specified in the `RandomForestClassifier` function in Python.

Gradient Boosting Classifier (GBC) was set with 50, 100, and 200 boosting stages (`n_estimators`), and a learning rate of 0.01, 0.1, 0.2.

Multilayer Perceptron (MLP) was implemented with hidden layers of sizes (100,), and (50, 50), a maximum of 1000 iterations (`max_iter`),alpha of 0.0001 and 0.001, and the ReLU and tanh activation function.

All models were implemented using the scikit-learn package (Pedregosa et al., 2011).

#### 2 Performance of the species-drug specific models

We analyzed the model's performance across various species-drug combinations using predictions from Set 2 to evaluate its effectiveness in different contexts. The model performed best on *Candida*

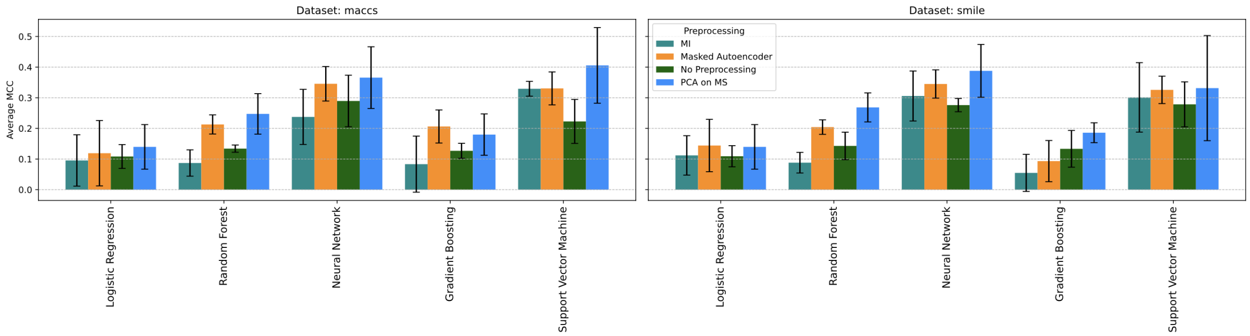


**Fig. 4:** MCC across pre-processing techniques and models for all species-drug pairs. Error bars show the standard deviation computed via the 3-fold cross validation.

*parapsilosis*, achieving an average MCC of 0.84 across drugs, followed by *Saccharomyces cerevisiae* (MCC 0.83) and *Candida albicans* (MCC 0.29). Similarly, among antifungals, the model performed best with micafungin (average MCC 0.89 across species), followed by 5-fluorocytosine (MCC 0.34) and caspofungin (MCC 0.33). Note that for five *Candida* species, the models were not directly trained on caspofungin. Instead, following EUCAST guidelines, isolates that were susceptible to anidulafungin as well as micafungin were considered susceptible to caspofungin. Overall, these findings highlight the model's ability to generalize across a diverse range of species and drugs. However, we observed significant performance variability across species-drug pairs and even between different drugs for the same species. For instance, while the model achieved strong results for *Candida albicans* with micafungin, it performed poorly when paired with posaconazole or voriconazole. The top-performing species-drug pairs in Set 2 are *Saccharomyces cerevisiae*–itraconazole, which achieved perfect predictions, followed by *Candida albicans*–micafungin (MCC 0.89), *Candida parapsilosis*–fluconazole (MCC 0.84), *Candida tropicalis*–5-fluorocytosine (MCC 0.68)*Saccharomyces cerevisiae*–fluconazole (MCC 0.65).

| Species | Drug | AUPRC | MCC | Balanced acc | Imbalance | TP | TN | FP | FN |
|---|---|---|---|---|---|---|---|---|---|
| saccharomyces cerevisiae | itraconazole | 1.00 | 1.00 | 1.00 | 0.63 | 5 | 3 | 0 | 0 |
| Candida albicans | micafungin | 0.86 | 0.89 | 0.90 | 0.06 | 4 | 85 | 0 | 1 |
| Candida parapsilosis | fluconazole | 1.00 | 0.84 | 0.97 | 0.15 | 3 | 16 | 1 | 0 |
| Candida tropicalis | 5-fluorocytosine | 0.58 | 0.68 | 0.75 | 0.13 | 1 | 13 | 0 | 1 |
| saccharomyces cerevisiae | fluconazole | 0.82 | 0.66 | 0.75 | 0.75 | 6 | 1 | 1 | 0 |
| Candida tropicalis | posaconazole | 0.98 | 0.48 | 0.85 | 0.87 | 9 | 2 | 0 | 4 |
| Candida albicans | anidulafungin | 0.60 | 0.40 | 0.69 | 0.26 | 12 | 56 | 9 | 11 |
| Candida albicans | itraconazole | 0.42 | 0.35 | 0.68 | 0.13 | 5 | 69 | 7 | 6 |
| Candida albicans | itraconazole | 0.97 | 0.14 | 0.62 | 0.92 | 35 | 2 | 2 | 13 |
| Candida glabrata | itraconazole | 0.84 | 0.04 | 0.52 | 0.73 | 6 | 2 | 2 | 5 |

**Table 1:** Performance metrics for the top-10 species-drug pairs, sorted by MCC. For caspofungin, susceptibility is inferred using both anidulafungin and micafungin (logical AND).

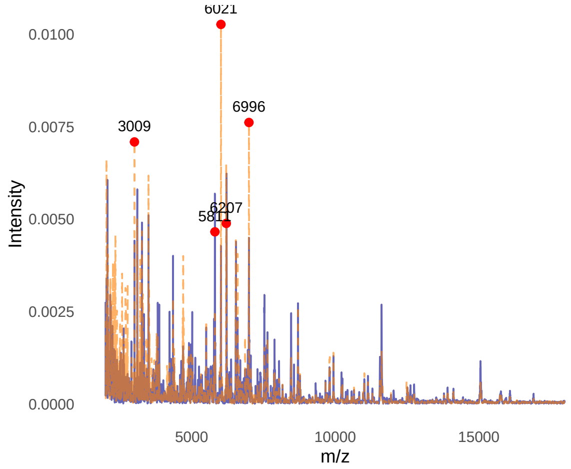


**(a)** *Saccharomyces cerevisiae - itraconazole and fluconazole*

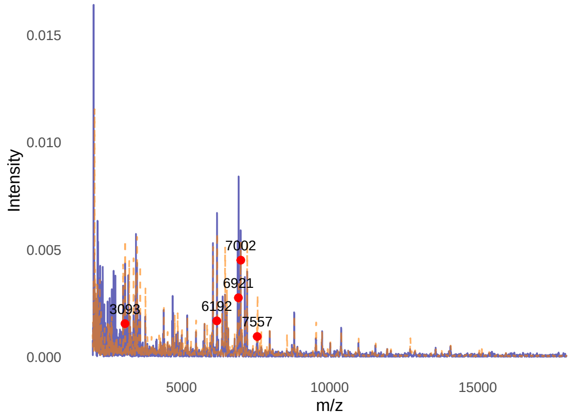


**(b)** *Candida tropicalis- posaconazole*

**Fig. 5:** Average mass spectra for susceptible (blue) and resistant (dashed & orange) pathogens. The peaks contributing the most to AMR prediction are annotated with red dots. (a) The min peak height value is $5.5 \times 10^{-4}$. The SHAP values were computed from 8 correctly classified observations. The 5 top mass spectra features differentiating between cases and controls are 3009, 5811, 6021, 6207 and 6996 m/z (±1). (b) The min peak height value is $4.7 \times 10^{-4}$. The SHAP values were computed from 10 correctly classified observations. The 5 top mass spectra peaks differentiating between cases and controls are 2092, 6192, 6921, 7002, 7557 m/z (±1).
